# Phenotype-driven screening reveals a causal role for the cortex in pupil control

**DOI:** 10.64898/2026.03.17.712501

**Authors:** Shingo Nakazawa, Yukako Tohsato, Tatsumi Hirata

## Abstract

Phenotype-driven forward screening offers a powerful strategy to discover neuronal substrates underlying physical and behavioral traits without prior anatomical or functional assumptions. Historically, such approaches have identified many important neuronal circuits using invertebrate model organisms, but application to mammals has remained limited by the lack of appropriate strategies. Here we quantitatively profiled 56 neurological phenotypic features across more than 200 adult mice when neurodevelopmentally classified neurons were activated or inhibited by chemogenetics. This screen yielded a wide spectrum of robust neurological phenotypes. Upon analyzing these phenotypes computationally, we narrowed down to the hypothesis that activation of cortical neurons enlarges pupil size. Experimental evidence using optogenetics and in utero electroporation indicated that this hypothesis is true. Our results provide proof of principle that phenotype-driven forward approach in mammals is a powerful alternative as a laboratory approach to uncover brain substrates, and offers a general framework for systematically mapping neural circuits that regulate physical and behavioral phenotypes.

## Introduction

In developmental neurobiology, neurons are fate-determined by the timing at which they differentiate and the spatial domain from which they differentiate^1^, and according to the fates, they make connections with specific partners to construct the functional brain^2^. Therefore, adult brain functions are ultimately shaped by neurodevelopmental principles. However, despite its conceptual importance, the causal contribution of neurodevelopmental origin to adult brain function remains largely unexplored, because of the lack of experimental strategies.

Neurogenic tagging provides a unique solution to this challenge^3,4^. By using tamoxifen (TM)-inducible Cre recombinase (CreER) driven by neurogenic transcription factors, neurons are permanently tagged at the time when they differentiate. This approach therefore allows genetic access to neuronal populations defined by their birth timing and developmental origin in the mature brain. Previously, we generated four neurogenic tagging transgenic mouse lines using the potential promoters of Neurog1, Neurog2, NeuroD1 and NeuroD4 genes to express CreER in the time window of neuron differentiation^4^ (Fig. 1A). Subsequently, administration of TM once during the neuronal differentiation time window irreversibly recombines genomic loxP sequences in the neurons that differentiate at similar timing from the same progenitor domain. (Hereafter, TM stage indicates the mouse embryonic stage at which TM was administered). The neurons tagged by this method are showcased in the NeuroGT database^4^ (https://ssbd.riken.jp/neurogt/)

**Figure 1.**
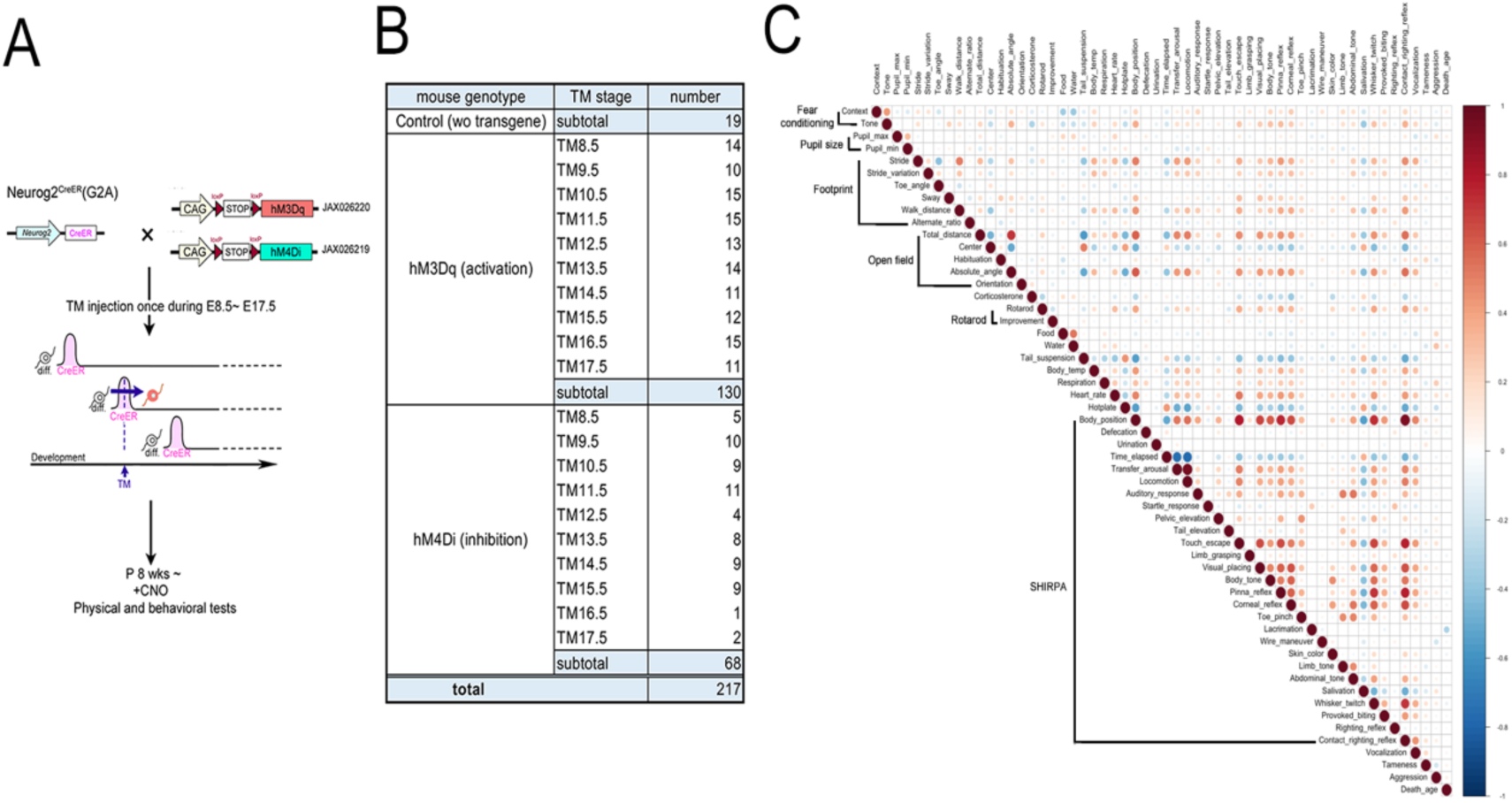
Neurogenic tagging-based forward phenotype screening. **A:** Schematic of the phenotype-driven forward screening strategy. Neurogenic tagging Neurog2^CreER^ (G2A) mice were crossed with either of an activation form (hM3Dq) or inhibition form (hM4Di) of DREADD mice. TM injection on a single day during E8.5–17.5 induces *loxP* recombination only in the cells expressing CreER. The resulting male mouse individuals were raised up to postnatal 8 weeks, intraperitoneally administered a DREADD ligand CNO, and their physical and behavioral phenotypes were assayed under the influence of CNO. **B:** The numbers of mice, and their genotypes and TM stages analyzed in the present screening. **C:** Pairwise correlations of 56 distinct phenotypic features. Spearman’s correlation coefficients were calculated using all the phenotypic scores.

In the present study, we employed the neurogenic tagging-based forward screening to systematically analyze physical and behavioral phenotypes of adult mice following causal manipulation of neurons classified by neurodevelopmental principles (Fig. 1A). Among the neurogenic tagging lines, the Neurog2^CreER^ (G2A) driver line labels exceptionally broad brain regions^4^, making it suitable for the initial phenotype-driven screening. Therefore, using this driver line, we selectively expressed either an activation form (hM3Dq) or inhibition form (hM4Di) of DREADD proteins^5^ in neurogenic-tagged neuronal populations (Fig. 1A). After the mice postnatally grew up till 8 weeks, an artificial ligand of DREADD, Clozapine N-oxide (CNO) was intraperitoneally administered to the mouse individuals, and their physical and behavioral phenotypes were assayed by a comprehensive battery of tests under the influence of CNO (Fig. 1A). Importantly, neither brain regions nor phenotypic outputs were predefined, enabling a phenotype-driven, unbiased forward screen. In this way, we aimed to identify brain substrates underlying physical and behavioral functions without prior assumptions.

## Results

### Phenotype-driven screening of neurodevelopmentally defined neurons

To systematically explore how neurons defined by neurodevelopmental principles contribute to adult brain function, we performed a phenotype-driven forward screening using neurogenic tagging mice. In total, 217 mice, including 19 control animals lacking any transgenes were subjected to a comprehensive battery of physical and behavioral tests (Fig. 1B). As detailed in the method, this screen incorporated multiple classes of phenotypic measurements. Several assays yielded fully quantitative readouts expressed in SI units (International System) such as body temperature or water consumption. Other assessments were adopted from standardized battery tests, including SHIRPA^6^ for mice, and scored on observational binary or ordinal scales. In the footprint, open field or rotarod tests, salient features were extracted and converted into numerical values. Altogether, 56 distinct phenotypic features were quantified and analyzed (Fig. 1C). These features were selected to broadly cover various neurological features such as motor, sensory or psychological conditions, thereby enabling a wide survey of phenotypic space. Although some features exhibited significant correlations (Fig. 1C), this comprehensive approach ensured maximal coverage of diverse physical and behavioral outcomes.

Chemogenetic manipulation of tagged neurons by DREADD proteins produced substantial phenotypic variabilities across TM stages (Supplementary Fig. 1). In general, neuronal activation using the excitatory DREADD hM3Dq elicited more pronounced effects than neuronal inhibition using hM4Di (Fig. 2, Supplementary Fig. 2, 3), consistent with the notion that activation of small neuronal ensembles can strongly perturb network activities. Notably, activation and inactivation did not typically produce mirror-opposite results, suggesting that many neural circuits operate through non-reciprocal mechanisms to regulate homeostatic outputs. Importantly, the magnitude of phenotypes observed in this forward screening was substantially greater than that reported in previous gene knockout–based screening studies^7–9^, likely reflecting the direct manipulation of neuronal activities rather than gene disruption.

**Figure 2.**
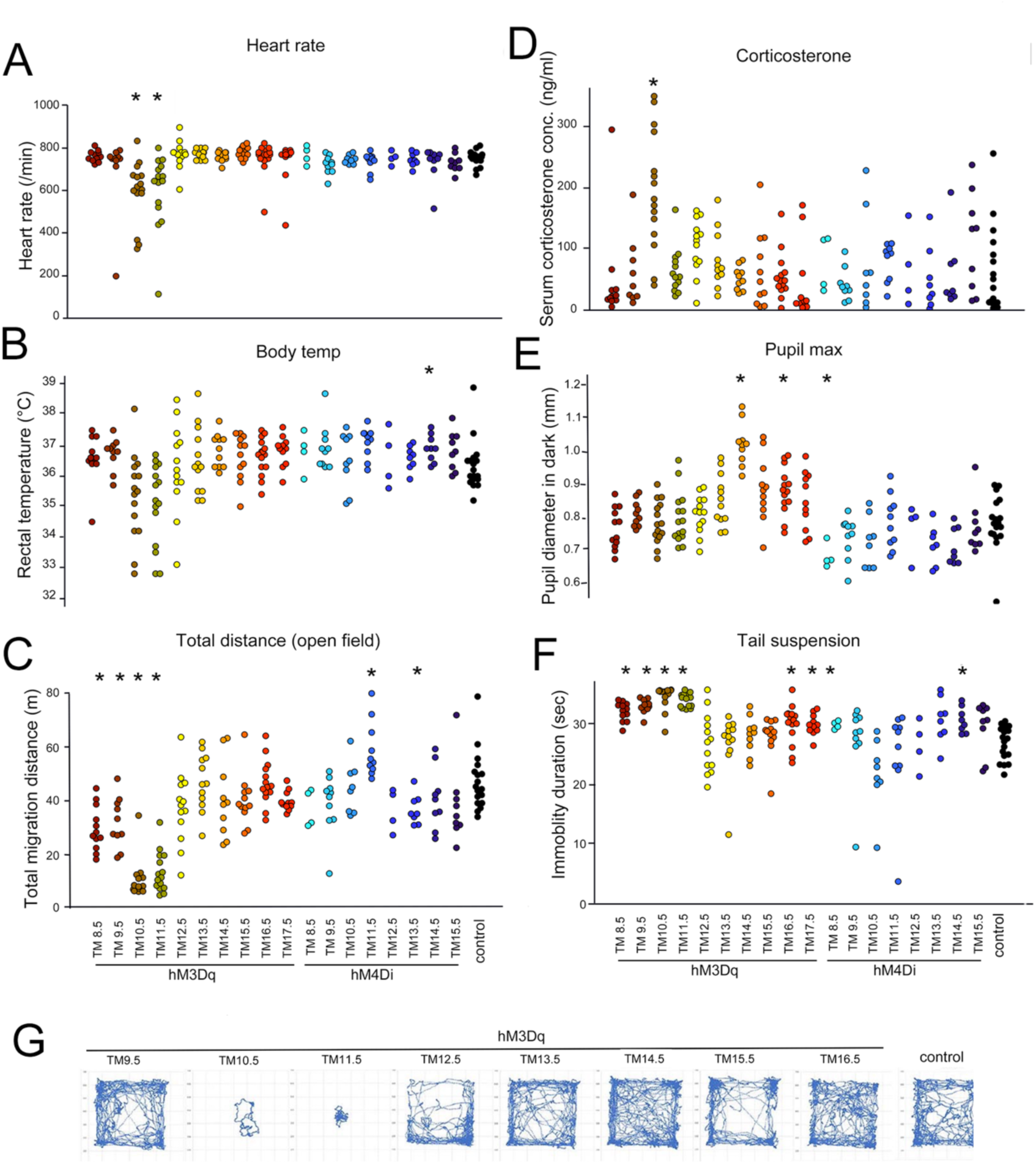
Neurological phenotypes revealed by physical and behavioral tests. **A-F:** Representative phenotypic features measured in quantitative scales of mouse individuals following chemogenetic activation or inhibition of neurogenic–tagged neurons. Each dot represents the phenotypic score of a single mouse. Colors indicate groups divided by TM stages shown on the bottom. Statistical significance (*) was assessed against the control group using Brunner-Munzel test with Bonferroni correction (*P* < 0.05). **G:** Representative migration trails of mice in the open field box for 10 min following the activation of neurogenic-tagged neurons.

### TM stage–specific phenotypes emerge from the screen

Closer inspection of individual phenotypes revealed distinct TM stage–specific effects. Activation of neurons tagged at TM10.5 and TM11.5 resulted in marked reductions in autonomic parameters, including heart rate, body temperature and respiration rate (Fig. 2A, B, Supplementary Fig. 2). Consistently, these mice exhibited profound behavioral suppression in the open field (Fig. 2C and G) and other tests (Supplementary Fig. 2, 3), remaining largely sedentary despite retaining some ability to perform on the rotarod (Supplementary Fig. 2). The serum stress hormone, corticosterone, dramatically increased when TM11.5 neurons were activated (Fig. 2D), displaying a clear difference from activation of TM10.5 neurons.

Among the phenotypes detected in the screening, pupil size exhibited a striking TM–stage dependence; activation of neurons tagged at TM14.5 or later produced robust enlargement of pupil size in darkness (Fig.2E). This phenotype was highly TM stage–specific and was not observed following activation of earlier tagged neuronal populations, suggesting the involvement of a distinct neuronal population. In contrast, other phenotypes such as immobilized time in the tail suspension test as an indicator of depression significantly increased following manipulation of neurons tagged at broad stages (Fig. 2F), potentially reflecting the involvement of multiple brain regions.

### Informatics analysis links pupil enlargement to cortical neurons

To visualize the global structure of the multidimensional phenotypic data, individual mice were embedded into a low–dimensional Uniform Manifold Approximation and Projection (UMAP)^10^ space using all quantified scores (Fig. 3A). Although mice corresponding to different TM stages partially overlapped in the UMAP space, their distributions appeared to slightly differ among TM stages. The most clearly separated cluster comprised mice in which TM10.5 and TM11.5 neurons were activated, showing pronounced sedentary behavior and autonomic dysfunction phenotypes.

**Figure 3.**
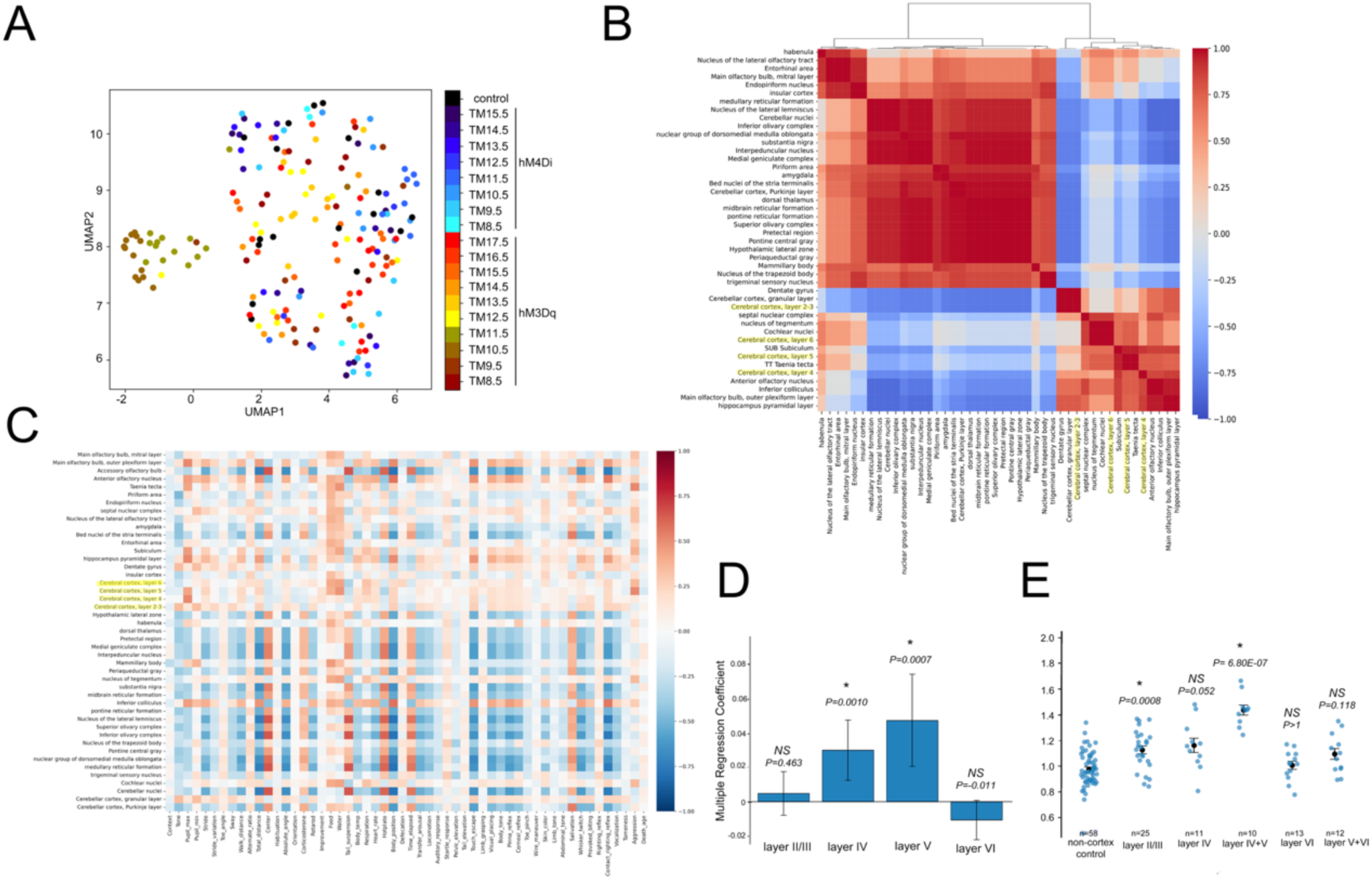
Informatics analyses. **A:** UMAP embedding^10^ of individual mice based on all 56 quantified phenotypic features. Each point represents one mouse, colored by TM stages. **B:** Heatmap of Pearson correlation coefficients between brain regions. Hierarchical clustering of brain regions was based on the inter-regional similarity using Ward’s method. Brain regions with similar phenotypic correlation profiles are positioned proximally. Cortical region names are colored in yellow. **C:** Heatmap of correlations between phenotypic scores and ontology-based brain regions. Cortical region names are colored in yellow. **D:** Multiple regression coefficients predicting the pupil size from cortical layers. Vertical lines indicate coefficients ± 95% confidence intervals. Sample size (n) is 120 in the analysis. *P* values are indicated in the panel. **E:** Strip plot of the pupil size upon activation of cortical layer and non-cortical control neurons. Black marks and lines show mean ± SEM. Statistical significances versus control group were calculated using Welch’s *t-*tests with Bonferroni correction. Sample sizes (n) and *P* values are indicated in the panel.

Because the Neurog2-CreER (G2A) driver line labels neurons across exceptionally broad brain territories^4^, multiple brain regions were simultaneously manipulated in each experimental condition (Fig. 3B). Consequently, direct attribution of specific phenotypes to individual brain regions was inherently difficult. To address this limitation, we focused on the cerebral cortex, which is robustly and selectively tagged by the Neurog2-CreER (G2A) line in a temporally ordered pattern^4^. Previous work demonstrated that cortical neurons are labeled sequentially from deep to superficial layers across TM stages^4^ in this line, providing a structured and interpretable anatomical framework. Using ontology-based anatomical annotations from the NeuroGT database, we computed correlations between tagged brain regions and phenotypic scores in hM3Dq and control groups (Fig. 3C).

When analyses were restricted to the cerebral cortex, a striking relationship with pupil size emerged (Fig. 3C). As described before, pupil size measured in darkness (Pupil_max) was significantly enlarged following activation of neurons tagged at TM14.5 (Fig. 2E). Previous work has shown that neurogenic tagging by the Neurog2-CreER (G2A) line is markedly reduced in non-cortical regions by this stage, leaving cortical layer IV neurons as the predominant labeled population^4^ (see also the NeuroGT database). Importantly, pupil enlargement induced by neuronal activation was reversibly suppressed by strong illumination to the eye (Pupil_min, Supplementary Fig.2), indicating that the pupillary light reflex remained intact.

To further quantify layer contributions, we performed multiple regression analysis using the tagging status of individual cortical layers shown in the NeuroGT database as predictors. To simplify the analysis, we used pupil sizes of hM3Dq and control groups as phenotypic outcomes (Fig. 2E). The regression analysis revealed significant positive contributions of cortical layers IV and V (Fig. 3D). Consistently, Welch’s *t*-tests against non-cortical controls showed significant pupil enlargement upon activation of cortical layers (Fig. 3E).

Together, these analyses indicate that activation of middle-to-superficial cortical layers is associated with pupil enlargement.

### Hypothesis testing: cortical activation is sufficient to induce pupil enlargement

We next directly tested the hypothesis that cortical activation induces pupil enlargement by two independent experimental approaches. First, cortical neurons tagged at TM14.5 were optogenetically activated using channelrhodopsin-2 (ChR2, Fig. 4A). To achieve localized stimulation, optic fiber guides were surgically implanted over the cortical skull either bilaterally at the center or unilaterally on the contralateral or ipsilateral side several days prior to recording. Subsequently, pupil size was continuously monitored during focal light stimulation. Optical activation induced a gradual and sustained pupil enlargement, reaching a maximum approximately 2–3 minutes after stimulation onset (Fig. 4B). This effect was robust and independent of the cortical site of stimulation (Fig. 4C); we observed no statistical significance in the effect among the center, contralateral and ipsilateral cortical stimulations. Mice were sacrificed after stimulation, and neuronal activation was verified histologically (Fig. 4D). Although dense dendritic labeling precluded precise laminar visualization of ChR2, robust neuronal activation was confirmed by expression of cFOS, an immediate early gene marker for neuronal activity. The cFOS expression suggested that initial activation of neurons in layer IV propagated vertically across cortical layers, with limited long-range spread laterally to the contralateral hemisphere (Fig. 4E). We did not detect a significant correlation between the number of cFOS-positive cortical neurons and the magnitude of pupil enlargement (Supplementary Fig 4A).

**Figure 4.**
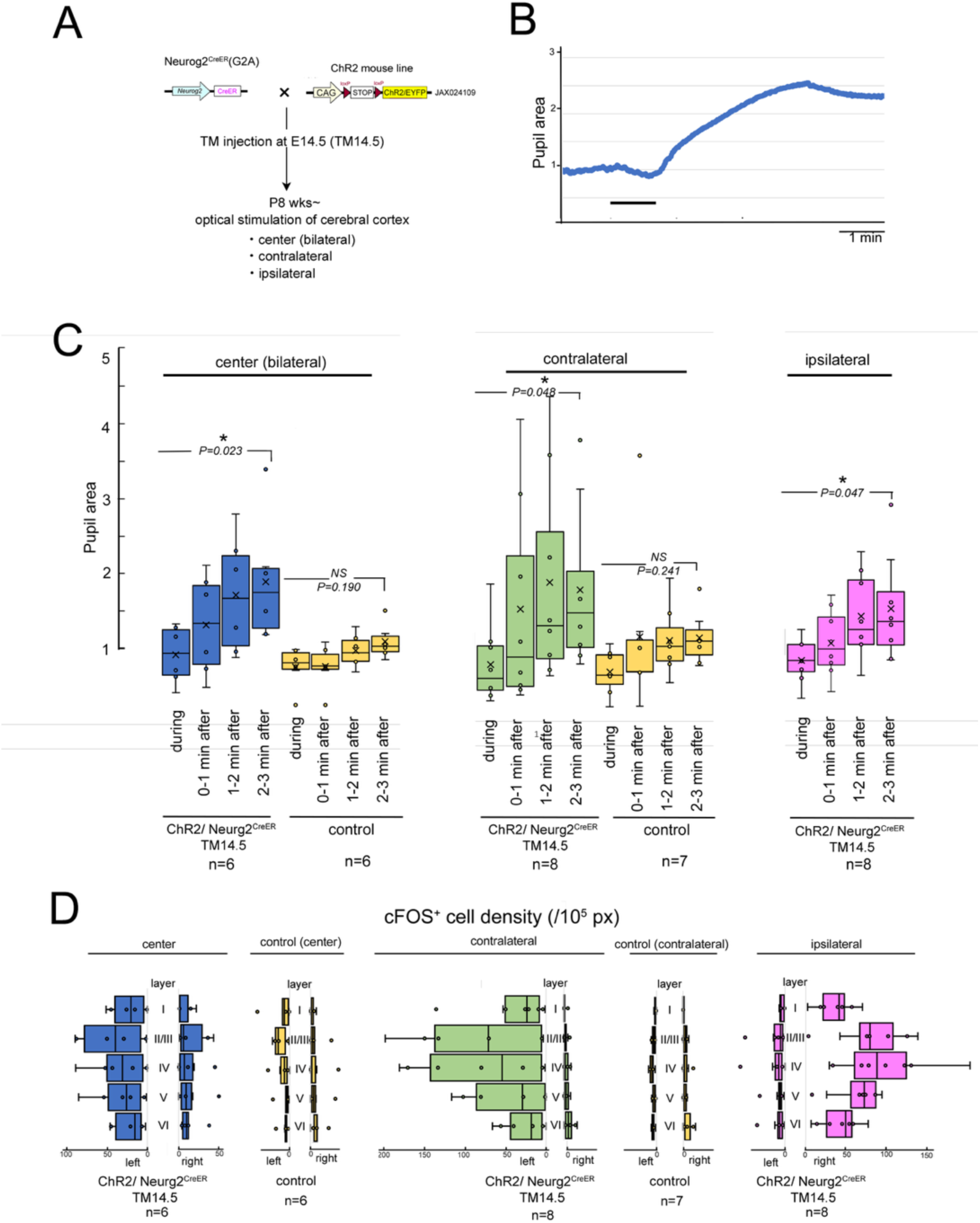
Optogenetic activation of cortical neurons. **A:** Experimental design for optogenetic activation of neurogenic tagged cortical neurons. Neurogenic tagging Neurog2^CreER^ (G2A) mice were crossed with ChR2 mice (Ai32), and TM was injected at E14.5. The resulting male mice were raised up to postnatal 8 weeks, and optical stimulation was given bilaterally at the center or unilaterally on the contralateral or ipsilateral side. **B:** Time course of pupil area in the right eye. The pupil area was expressed as a relative value to the average area of the same mouse before optical stimulation. The horizontal bar indicates the time interval of optical stimulation. **C:** Boxplot showing relative pupil area of right eyes after optical stimulation over the center, contralateral or ipsilateral cortex. X marks the mean. The pupil area was slightly smaller during the stimulation, possibly due to the unavoidable leakage of light over the eye, but soon enlarged in ChR2-expressing groups regardless of the position of stimulation. As the control group, C57BL/6 wild mice without any transgene were used. *P* values were calculated by paired *t*-tests between before and 2-3 min after the stimulation. The genotypes and the numbers of mice analyzed are shown at the bottom. **D**_3_: _0_Distribution of cFOS-positive neurons in layers of left and right somatosensory cortices. The same mice in **C** were analyzed at postmortem.

As a second complementary approach, we performed cortex-specific chemogenetic activation using the excitatory DREADD, hM3Dq (Fig. 5A). A plasmid encoding hM3Dq fused with mCherry under a ubiquitous promoter was introduced unilaterally into the dorsolateral cortex of wild-type embryos on a random side by in utero electroporation. Although this procedure was only performed on ICR mice yielding pups with albino eyes, we were able to measure pupil sizes after the animals matured to adulthood, allowing within-animal comparisons following administration of CNO or vehicle (PBS) (Fig. 5B). Consistent with previous reports^11^, electroporation at E12.5–E15.5 resulted in layer-specific expression of hM3Dq in an inside-out gradient depending on developmental timing (Fig. 5C). Administration of CNO induced significant pupil enlargement in mice electroporated at E14.5 and E15.5 (Fig. 5B), in which hM3Dq was expressed in superficial cortical layers (Fig. 5C). Electroporation at E12.5 and E13.5, which primarily targets deep cortical layers, did not produce statistically significant pupil enlargement (Fig. 5B). Although mice expressed hM3Dq only unilaterally either in the left or right cortical hemisphere on the electroporated side (Fig. 5C), the pupil enlargement was not different between right and left eyes or between contralateral and ipsilateral eyes to the electroporated side (Supplementary Fig. 5). The cFOS analysis indicated that activation of cortical neurons by hM3Dq propagated vertically and partially to the contralateral cortical side (Fig. 5D). Again, the correlation between the number of cFOS-positive neurons and pupil enlargement was not significant (Supplementary Fig. 4B).

**Figure 5.**
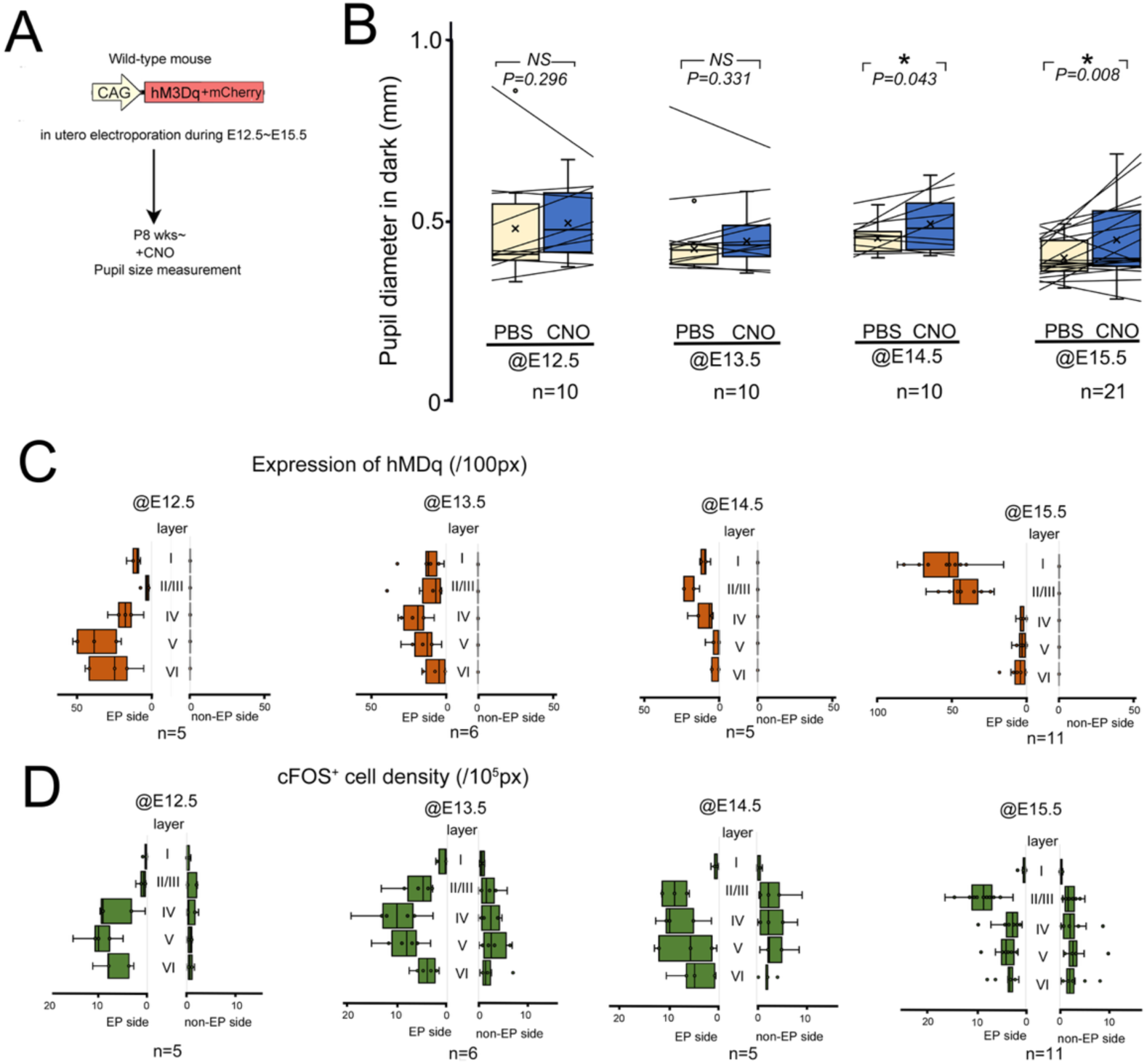
Chemogenetic activation of cortical neurons after restricted introduction of the activation form of DREADD. **A:** Experimental design for cortex-restricted chemogenetic activation. An expression plasmid for hM3Dq-mCherry was in utero electroporated unilaterally to the dorsolateral cortex of ICR wild-type embryos at E12.5-15.5. After the resulting male mice grew up till postnatal 8 week, CNO or vehicle PBS was intraperitoneally administered to the mice, and the pupil sizes of both eyes were measured in a dark room. **B:** Boxplot showing the pupil diameters from eyes at 20-50 mins after administration of PBS (pale yellow) and CNO (blue) in ICR mice whose cortices were in utero electroporated for hM3Dq at E12.5, E13.5, E14.5 and E15.5. The black line shows the difference of pupil diameters in individual eyes between PBS and CNO treatments. X marks the mean. One-tailed paired t-tests were performed between PBS and CNO treatments. **C, D:** Distribution of hM3Dq-positive areas and FOS-positive neurons in cortical layers of somatosensory areas on the in utero electroporated (EP) and unelectroporated (non-EP) sides. The same mice in **B** were analyzed at postmortem.

## DISCUSSION

In this study, we systematically examined physical and behavioral phenotypes in adult mice by manipulating the activity of neurons classified according to their neurodevelopmental origin. Using an informatics-based approach, we derived the hypothesis that activation of cortical neurons induces pupil enlargement, and we experimentally demonstrated that this hypothesis is correct. This result provides direct evidence that a phenotype-driven strategy can successfully identify functionally relevant neuronal circuits without prior anatomical or functional assumptions. Although the present study prioritized cortical analyses, our ongoing high-resolution non-cortical mapping using additional neurogenic tagging lines with more restricted spatial tagging is expected to further uncover neuronal circuits that regulate other physical and behavioral phenotypes.

Pupil size is widely used as an indicator of arousal state in animals^12,13^ and is primarily regulated by the noradrenergic system originating from the locus coeruleus^14^. Cortical activity has been proposed to be a component of the dynamic arousal system^15^. While previous studies have mainly reported correlations between cortical activity and pupil size^16–18^, the present study demonstrates a causal relationship, showing that direct activation of cortical neurons induces pupil enlargement. We observed a minor discrepancy in the peak timing of pupil enlargement between neurogenic tagging and in utero electroporation experiments, which may reflect differences in neuronal subtypes labeled by these two methods. Nonetheless, the present phenotype-driven screening approach identified cortical neuronal circuits as significant contributors to pupil enlargement.

With the exception of a few well-characterized brain regions^4^, neurons that differentiate at similar developmental time points from the same progenitor domains are generally dispersed throughout the adult brain without clear correspondence to conventional anatomical landmarks (NeuroGT). Despite this apparent spatial dispersion, our phenotype-driven screening revealed TM–specific peaks that were uniquely associated with individual physical and behavioral phenotypes. These findings suggest that the neurodevelopmental principle indeed determines functional identity of neurons in the adult brain.

## Methods

### Mice

Neurogenic tagging mouse strains were generated as described previously^3,4^. Activation type DREADD mice^19^ {JAX#026220,Tg(CAG-CHRM3*,-mCitrine) *^Ute^*}, inhibition type DREADD mice^19^ {JAX#026219, *Gt(ROSA)26Sor^tm1(CAG-CHRM4*,-mCitrine)Ute^*} and optogenetics mice^20^ {JAX#024109, *Gt(ROSA)26Sor^tm32(CAG-COP4*H134R/EYFP)Hze^*} were obtained from the Jackson laboratory. C57BL/6 and ICR wild-type mice were purchased from SLC Inc. (Japan). All mice were maintained in the animal facilities of the National Institute of Genetics under controlled environmental conditions. The day a vaginal plug was detected was designated as embryonic day 0.5 (E0.5), and the day of birth was designated as postnatal day 0 (P0). TM administration was performed by intraperitoneal injection of 250 µL of corn oil (Sigma-Aldrich, Merck, MO) containing 9 mM tamoxifen (Sigma-Aldrich) and 5 mM progesterone (Fujifilm Wako, Japan) into a pregnant mouse at the appropriate gestational stage. Because TM injection often delays delivery, pups not born by E19.5 were delivered by Caesarean section and fostered by ICR surrogate mothers. All animal experiments were approved by the Animal Committee at the National Institute of Genetics and carried out according to the guidelines.

### Physical and behavioral tests

Neurog2^CreER^(G2A) heterozygous males were crossed with homozygous activation-type or inhibition-type DREADD females and treated as described above. Upon weaning of the pups, male mice were selected and genotyped from the tail genomic DNA for the CRE recombinase and DREADD alleles as described previously^3,4^ or on the Jackson laboratory website (https://www.jax.org/). The mice were individually housed thereafter. Once the male mice became 8 weeks of age, 0.02 mg of CNO (Cayman Chemical, MI) dissolved in PBS was injected intraperitoneally into each individual, and the mice were tested 20-90 mins after CNO injection except for food and water intakes and corticosterone assays. All tests were conducted between 9:00 and 16:00.

**Fear conditioning:** Conditioning was performed by pairing three tone stimuli (4 sec each, at 30 sec-intervals) with electric foot shocks (2 sec, 75V) in a shocking chamber^21^ (O’Hara & Co., Ltd., Japan) following CNO administration. On the following day, the conditioned mice without receiving CNO were first placed in the same chamber and recorded for five minutes without electric stimulation (Context). Subsequently, the mice were placed in a different white box (ST box 25, JSJ Astage, Japan) and exposed to the tone cue for 4 min without electric stimulation (Tone). Freezing behavior was quantified using DeepLabCut software^22^. **Pupil size:** Pupil images were acquired using a digital camera (Anyty, 3R Solution Corp., Japan) mounted on a dissecting microscope (SZ40 with TL3-100 illumination, Olympus, Japan). Measurements were obtained under the dark condition (∼1.8 lux; Pupil_max), and then the bright condition (∼9000 lux; Pupil_min) illuminated by an LED pen light (IYP365, Lumintop). The pupil diameter along the long axis was measured using Fiji/ImageJ^23^, and the diameters of right and left pupils were averaged. **Footprint**: Hind paws of the mice were dipped in black stamp ink (SGN_250K, Shachihata Inc., Japan), after which the mice were allowed to walk through a straight tunnel (7 × 9 × 47 cm) lined with white filter paper. From scanned foot prints, the total walking distance (walk_distance), averaged stride length (Stride), stride length variability (Stride_variation), averaged sway distance (Sway), toe angle (Toe_angle), and the ground-to-air contact time ratio (Alternate_ratio) were measured using Fiji/ImageJ^23^ and Microsoft Excel (2021 for Mac). **Open field:** Mice were placed in the center of a white square box (45 × 45 × 40 cm, LE825S, Panlab, Spain) and video-recorded from above for 10 min using a 1080P camera (W88S, Imilab, China). The videos were analyzed using DeepLabCut^22^, and total distance traveled (Total_distance), percentage of time spent in the 20 X 20 cm-center (Center), migration speed change (Habituation, = the last 2 min-distance divided by the first 2 min-distance), cumulative changes in the body axis (Absolute_angle) and overall direction of movement (Orientation) were calculated using Microsoft Excel (2021 for Mac). **Corticosterone:** Serum corticosterone levels were measured using a corticosterone ELISA kit (Cayman Chemical) from frozen serum samples collected at two hour after CNO injection. **Rotarod:** The rotarod apparatus (O’Hara & Co., Ltd.) accelerated from 2 to 40 rpm over 300 seconds following the previous method^21^. Latency to fall was measured 5 times at 150-second intervals, and averaged (Rotarod). Motor learning (Improvement) was calculated as follows: (the last latency*2 + the penultimate latency)/ (the first latency*2 + the second latency). **Food & Water:** Food and water intakes during the 0-5 hour period after CNO administration were measured by weighting initial and remaining food and water. **Tail suspension:** Mice were suspended by the tail with tape for 360 sec, and the immobile time within the time window was measured^24^. **Body temp:** Rectal temperature was measured using a microprobe thermometer (BAT-12, Physitemp, NJ). **Respiration_rate:** Movement of the mouse chest was zoom-in videotaped (Handycam, Sony, Japan) and respiration frequency (breaths/min) was analyzed using tanaMove (version1)^25,26^. **Heart_rate:** Heart beating sound was recorded using an IC recorder (Voice-Trek V+75, Olympus, Japan) and the frequency of heart beating (beats/min) was measured using Praat software^27^ (https://www.fon.hum.uva.nl/praat/). **Hotplate:** Mice were placed on a preheated hotplate (NHP-M30N. Nisshinrika, Japan) at 58℃, and the time until the mouse licked its paw or jumped away was measured, within a cutoff time of 60 sec. **SHIRPA:** Body_position, Defecation, Urination, Time_elapsed_before_move, Transfer arousal, Locomotion, Pelvic_elevation, Tail_elevation, Touch_escape, Limb_grasping, Visual_placing, Body_tone, Pinna_refex, Corneal_reflex, Toe_pinch, Lacrimation, Wire_maneuver, Skin_color, Limb_tone, Abdominal_tone, Salivation, Provoked biting, Right_reflex, and Contact_righting_reflex were assessed as described on the RIKEN BRC website^28^(https://ja.brc.riken.jp/lab/jmc/shirpa/) according to the uploaded video using the standard tools (O’Hara & Co., Ltd.). Auditory_response was scored separately from the Startle_response in this study. Whisker_twitch was evaluated according to the General Health and Neurological Screening protocol (GHNS) on the CBSN platform (https://cbsn.neuroinf.jp/database/item/88). **Vocalization, Tameness, Aggression, Death_age:** Vocalization and aggression were scored in the binary scale during the SHIRPA testing, whereas tameness^25^ was assessed similarly during the rotarod test. The dates of birth and death were recorded for mouse individuals, and the life times were calculated with a cutoff time of 100 days (Death_age).

### Optogenetics

Male mice from 8 weeks old onwards with appropriate genotypes were anesthetized with isoflurane (YS-18 AnesII, Biomachinery, Japan) and the skull was exposed and polished using a micrograinder (Twister200, Sea Force, Japan) with a diamond point (ED107, Crestetech, Japan). A 5-mm-diameter guide (SC050S, Iwata Manufacturing Co., Japan) of an optic fiber was affixed to the thinned skull at the center (bilateral), contralateral, or ipsilateral side using adhesive (LBR-005 Loctite, Henkel AG, Germany) and tightly sealed with a dental cement (Super-Bond C&B kit, Sun Medical, Japan). The guide is sufficiently large to cover the majority of cortical surface. After the mice recovered, they were kept in the home cage for 4-5 days. The mice were re-anesthetized with isoflurane and the head was fixed in a holder (SGM-4, Narishige, Japan) on a heating pad (RH-210, Marukan, Japan). The right eye was imaged using a CMOS camera (CS165MU, Thorlabs, NJ) mounted on a dissecting microscope (SZ-01, Olympus) under infrared LED illumination (LIU850, Thorlabs). An optical fiber (inner diameter 3 mmΦ, outer diameter 5 mmΦ, LLG03-4H, Thorlabs) connected to a blue LED (M470L5, Thorlabs) was inserted into the implanted guide. Light leakage was minimized using black tape (EL-12, 3M Japan) and sealing compound (Neoseal B3, Nitto Chemical, Japan). Output power was adjusted to 140 mW (20mW /mm^2^) at the fiber tip using an LED driver (LEDD1B, Thorlabs), and pulsed light (50ms off/ 50ms on) was created using an Arduino Uno (Arduino, Italy). Pupil area was quantified using the threshold function in Fiji/ImageJ^23^ and expressed as the relative size to that at 1 min before the optical stimulation.

### In utero electroporation

pAAV-PTRE-tight -hM3Dq-mCherry^29^ was obtained from Dr William Wisden (Addgene plasmid # 66795) and the coding sequences for hM3Dq fused with mCherry were subcloned into a CAGGS expression plasmid^30^. In utero electroporation was performed as described previously^31^. In brief, pregnant ICR wild-type mice were anesthetized with a combination of isoflurane (1.0% in air) and pentobarbital sodium (80 mg/kg body weight, Tokyo Chemical Industry, Japan). The uterus was exposed after abdominal incision, and ∼2 μl of plasmid solution was injected into either side of the telencephalic ventricle unilaterally. Five square electric pulses (30V at E12.5, 38V at E13.5, 40V at E14.5 and 43 or 44 V at E15.5, for 50-ms duration at 200-ms intervals) were delivered using forceps-type electrodes (CUY650P5 or CUY650P2, Nepa Gene, Japan) connected square-pulse generator (CUY21, BEX, Japan), targeting at the dorsolateral cortex. When the resulting male pups grew up till 8 weeks, sizes of pupils were bilaterally measured as described in the phenotypic test. The number of mice used in the measurement were 5 electroporated at E12.5 (3 in right and 2 in left hemispheres), 6 at E13.5 (3 in right and 3 in left hemispheres), 5 at E14.5 (3 in right and 2 in left hemispheres) and 11 (6 in right and 5 in left hemispheres) at E15.5.

### Immunohistochemistry

Frozen sections were prepared as described previously^32,33^. Sections were heat-treated in an antigen retrieval solution (HistoVT One, Nacalai Tesque, Japan) in an autoclave at 105°C for 2 min and washed with 10 mM Tris-HCl (pH 7.4), 130 mM NaCl and 0.1% Tween 20 (TBST). They were incubated with rabbit anti-cFOS monoclonal antibody (1:1000, ab222699, Abcam, UK) diluted in PBS containing 0.5% blocking reagent (FP1020, PerkinElmer, MA) for two days at 4°C, and the labeling was detected using donkey Alexa488-conjugated anti-rabbit IgG (1:1000, Life Technologies, CA). In utero electroporated samples were double-stained with rat anti-mCherry antibody (1:1000, M11217, Invitrogen, MA) and Cy3-conjugated AffiniPure donkey anti-rat IgG (1:1000, 712-165-153 Jackson Immunoresearch, PA). All the sections were counterstained with DAPI (4’,6-diamidino-2-phenylindole, 045-30361, CAS#28718-90-3, Fujifilm Wako, Japan).

### cFOS and hM3Dq expression analyses

Three sections of somatosensory cortices (SSCs) in right and left hemispheres from each mouse were immunostained. For each section, three optical sections were merged using CellSens FV software (Evident, Japan) under a confocal microscope (FV3000, Evident). The numbers of cFOS-positive cells were manually counted on the images using the Cell Counter plugin in Fiji/ Image J. The hM3Dq (mCherry)-expressing areas were automatically measured using the threshold function in Fiji/ Image J. Cortical layers were identified by DAPI staining and stored using the ROI manager tool.

### Statistical analysis

RStudio (2024.04.1+748)^34^ with R version 4.3.2 package^35^ was used for statistical analysis. Spearman’s rank correlations among 56 phenotypic features were calculated and visualized with corrplot, using pairwise complete observations to handle missing values (Fig. 1C). For quantitative phenotypic scores in the screen, group medians were compared using the Kruskal–Wallis test; categorical measurements were compared across groups using Fisher’s exact test (Supplementary Fig. 1). When significant differences were found, multiple comparisons against the control group were performed using Brunner-Munzel test with Bonferroni correction for quantitative measurements (Fig. 2, Supplementary Fig. 2) and Fisher’s exact test with Bonferroni correction for categorical measurements (Supplementary Fig. 3). The asterisks in the figures indicate *P* < 0.05, whereas no significance (NS) means *P* > 0.05.

Additional analyses were performed in Python^36^ (version 3.11.4) using NumPy^37^ (2.2.6), pandas^38^ (2.3.0), SciPy^39^ (1.15.3 or 1.17.1), statsmodels^40^ (0.14.4), scikit-learn^41^ (1.7.0), umap-learn^10^ (0.5.7), matplotlib^42^ (3.10.3), and seaborn^43^ (0.13.2). For global structure in the multidimensional phenotype space, UMAP^10^ was applied (Fig. 3A). Ontology-based tagged brain regions were selected for immunostaining of β-Gal-nls reporter using G2A line from NeuroGT database. To visualize similarity relationships among brain regions, a symmetric brain-region × brain-region correlation matrix was constructed using region-wise Pearson correlation coefficients (Fig. 3B). The resulting similarities were converted to distances, and hierarchical clustering was performed using Ward’s method. The clustered matrix was visualized as a heatmap with corresponding dendrograms (Fig. 3B). To relate phenotypes to anatomy, a binary brain-region × TM-stage matrix was first constructed (Fig. 3C), with 1 indicating the presence of a detected signal and 0 indicating its absence; controls were assigned 0 for all regions. Correlations between phenotypic scores and ontology-based brain regions were then computed using pairwise complete observations, after excluding zero-variance columns within the merged dataset. Layer contributions to the pupil size were evaluated by multiple regression using the annotation on cortical layers tagged in NeuroGT as predictors, including intercept terms (Fig. 3D). For group-wise pupil comparisons (Fig. 3E), datasets were stratified by combinations of layer-positive regions, and Welch’s *t*-test against controls were performed with Bonferroni correction (Fig. 3E). In the informatics analyses (Fig. 3), mouse individuals that contain missing values were excluded, and phenotypic features were min-max scaled prior to the analyses. In Fig. 3C-E, phenotypic datasets of hM3Dq and control groups were selectively used.

Through hypothesis testing, differences in the mean of pupil sizes were evaluated by one-tailed paired *t*-test using Microsoft Excel for Mac (Fig. 4, 5). Correlations between the number of cFOS-positive cells and pupil sizes were analyzed using CORREL function in the Microsoft Excel (Supplementary Fig. 4). Pupil diameter differences between right and left eyes and contralateral and ipsilateral eyes were compared using Welch *t*-test in Microsoft Excel for Mac (Supplementary Fig. 5). Sample sizes (n) and *P* values are indicated in the panel.

## Acknowledgements

The authors thank Jun-ichi Miyazaki for pCAGGS, William Wisden for pAAV-PTRE-tight-hM3Dq-mCherry, Akira Tanave for tanaMovie and Aki Masuda for excellent support in utero electroporation. We also thank Tsuyoshi Koide for lending various apparatuses and providing helpful comments in mouse behavioral analyses, Hiroshi Mori and Koichi Higashi for advice about informatics analyses, Takeo Katsuki for support to construct the optogenetic system and Yan Zhu for helpful support in utero electroporation and comments on experimental data and the manuscript. This work was supported by the grants from MEXT/JSPS KAKENHI Grants (23K27272) and IU-REAL Collaborative Research Grant to T. H.

## Author contributions

T.H. conceived the research, performed the experiments, analyzed the data, and wrote the manuscript with help from other authors. S.N and Y. T. performed informatics analyses.

## Materials & Correspondence

Correspondence and material requests should be addressed to T.H.

## Data availability

Row images and additional datasets generated during this study are available from the corresponding author (TH) upon reasonable request.

---

**Supplementary Figure 1.**
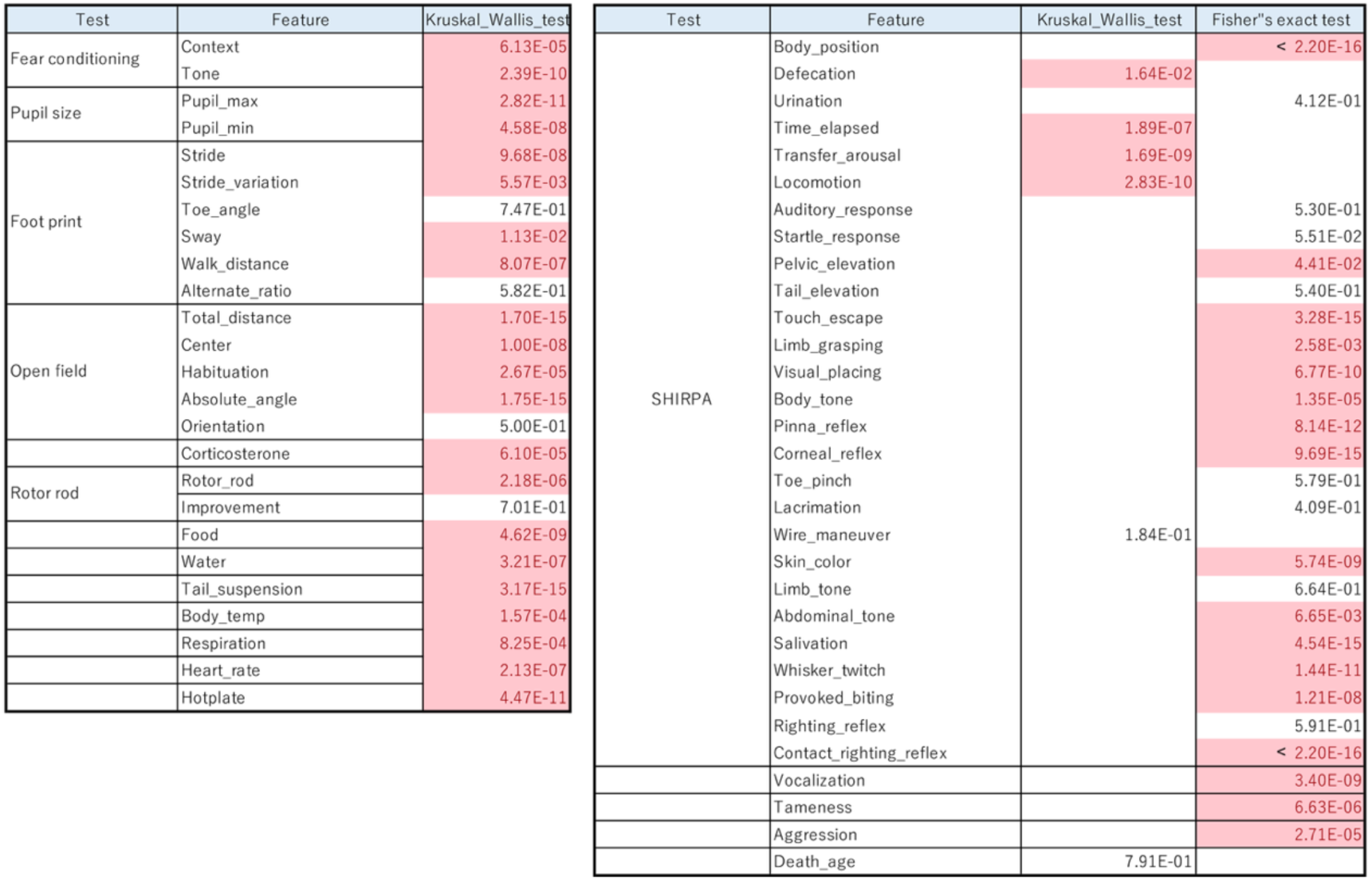
Statistical *P* values under the null hypothesis that median of phenotypic features are the same among TM and control groups. Kruskal–Wallis tests were conducted for quantitative measurements, whereas Fisher exact tests were conducted for categorical phenotypes in observation scales as long as computation was feasible. *P* < *0.05* were highlighted in red.

**Supplementary Figure 2.**
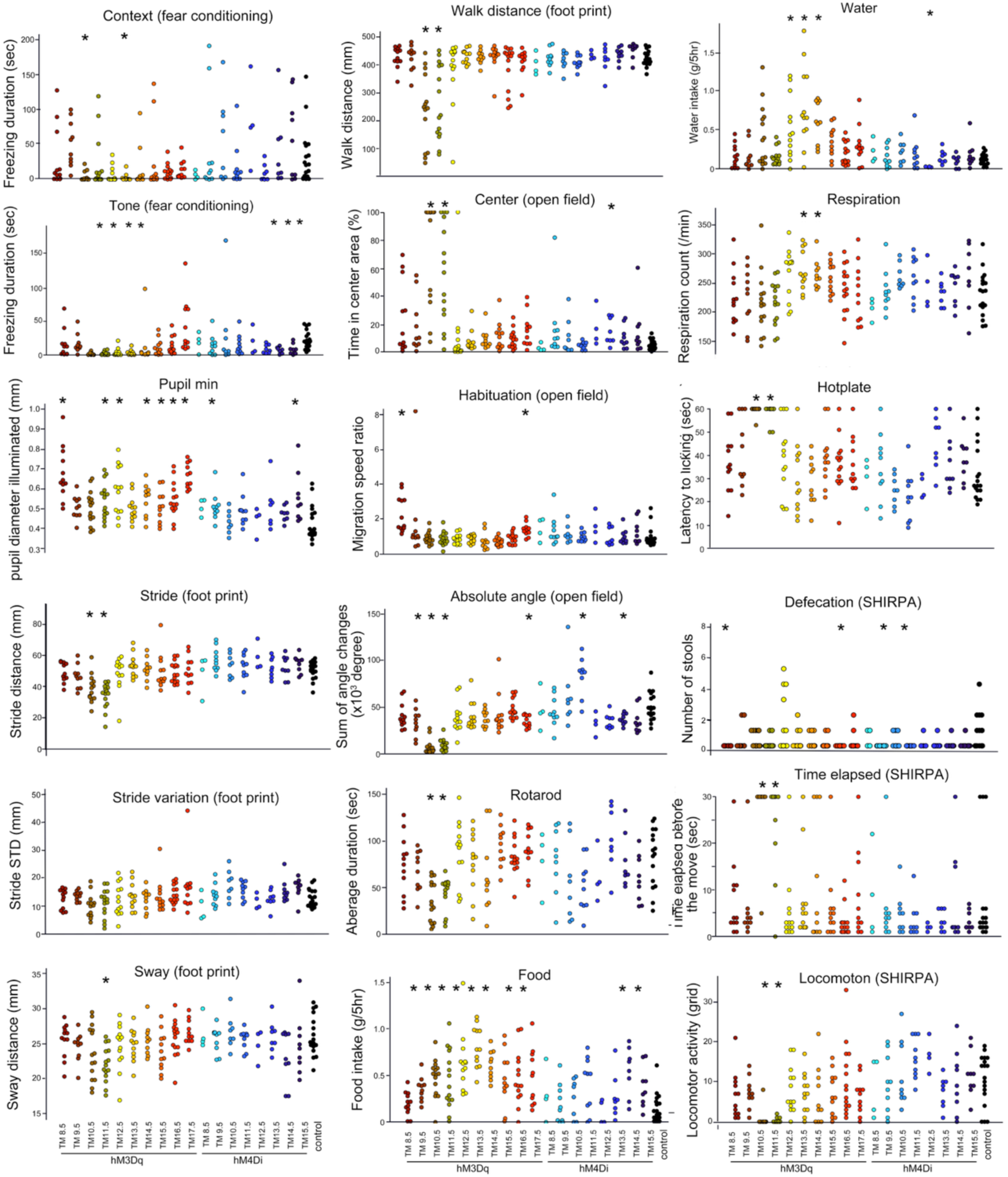
Quantitative phenotypic scores of individual mice after CNO administration. Significant features not addressed in Figure 2 are presented. Each colored dot indicates the value of a single mouse belonging to the group indicated at the bottom. *Asterisk shows statistical significance (*<0.05*) by Brunner-Munzel test with Bonferroni correction.

**Supplementary Figure 3.**
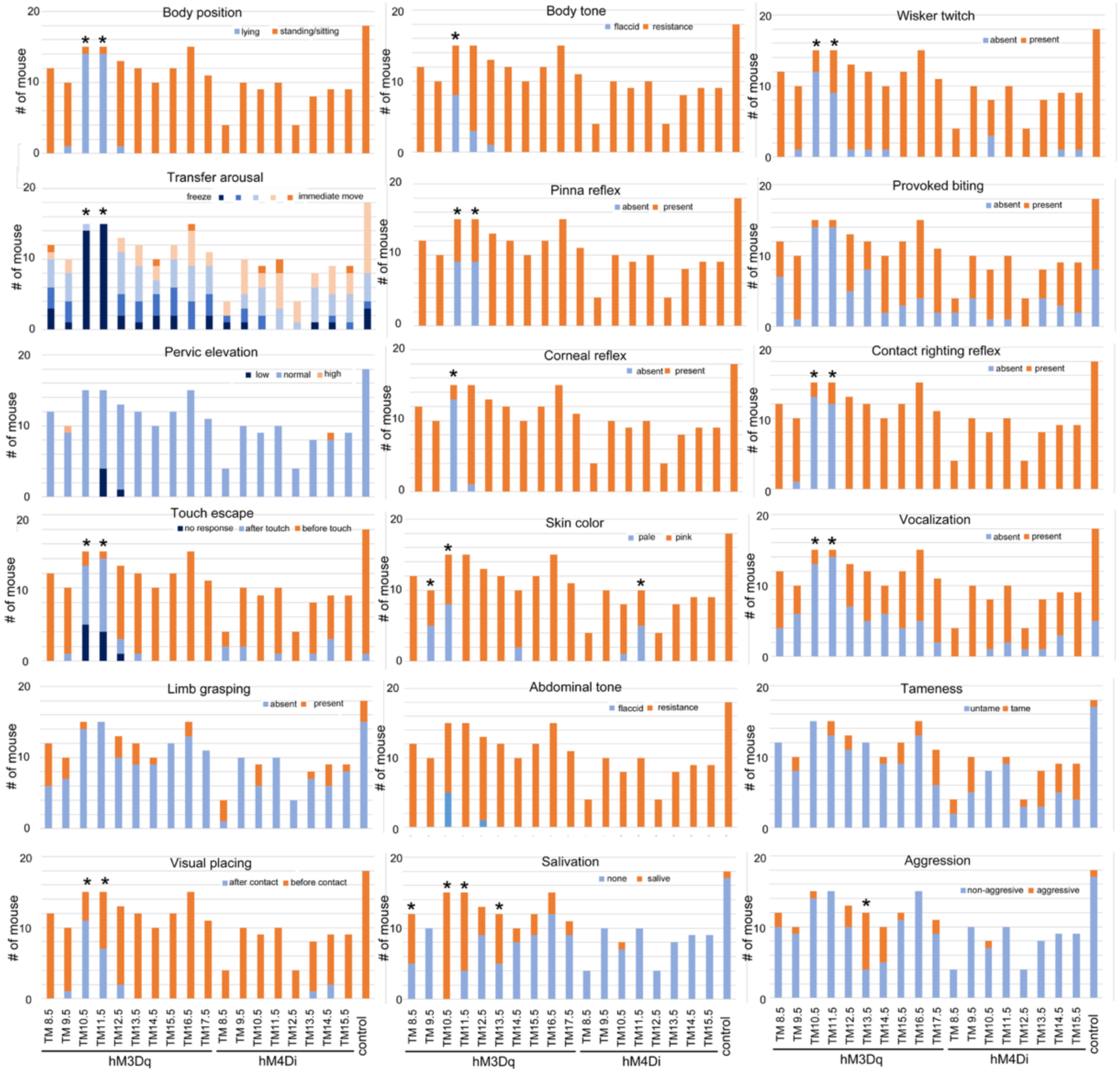
Phenotypic scores measured in observational scales after CNO administration. Bars indicate the number of mice showing the indicated phenotypes. *Asterisk indicates statistical significance (*<0.05*) when the phenotypic frequency in TM groups was compared with that in the control group by Fisher’s exact test with Bonferroni correction.

**Supplementary Figure 4.**
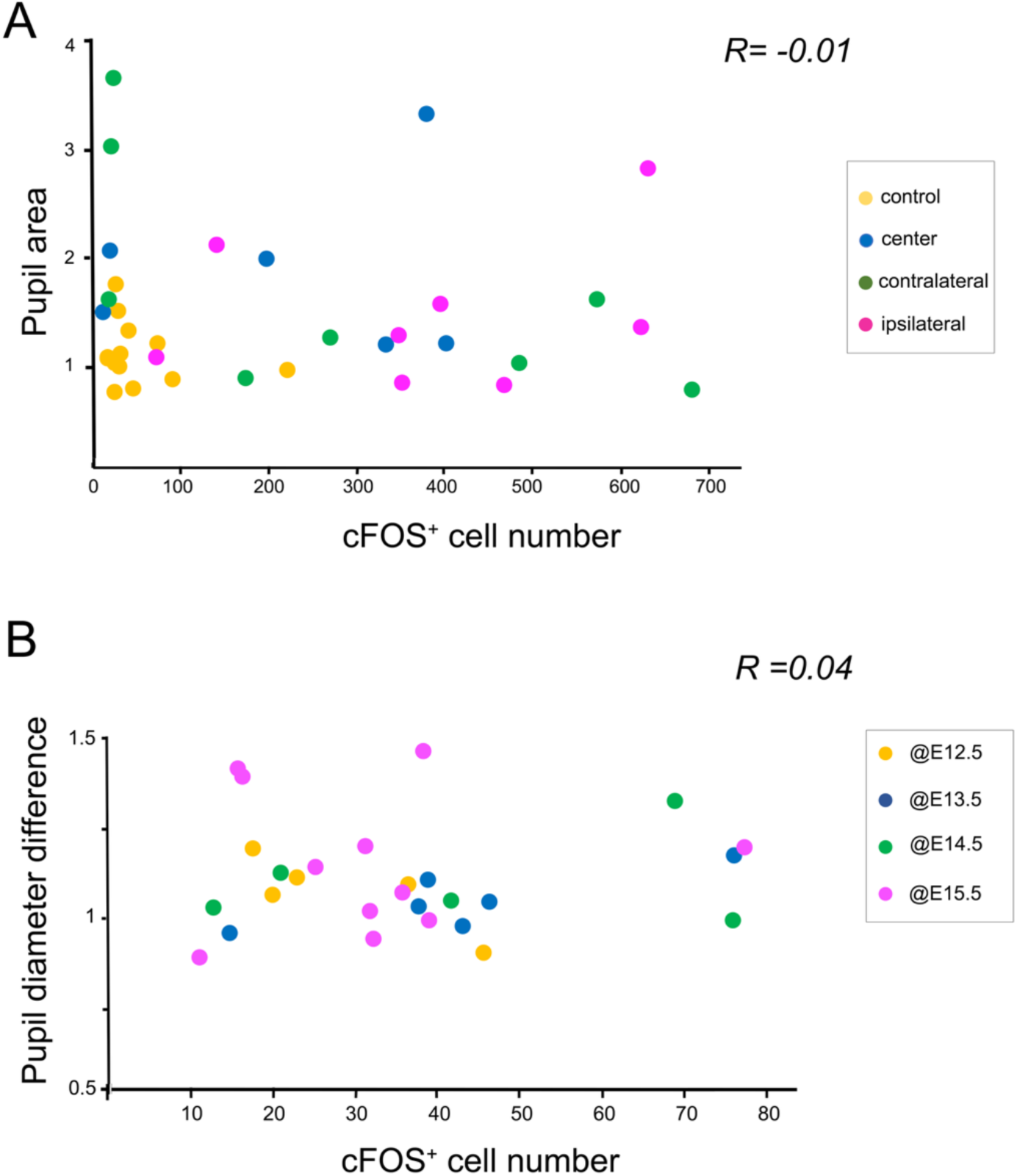
**A, B:** Correlations in the number of cFOS-positive neurons and the pupil size in optogenetics experiment (**A**) and cortex-specific introduction of hM3Dq by in utero electroporation (**B**). The number of cFOS-positive neurons are calculated as sum of all cortical layers in somatosensory areas on both sides. The y-axis in (**A**) shows the relative pupil area after optical stimulation as in Fig. 5, whereas that in (**B**) shows the pupil diameter differences between PBS and CNO treatments. The correlation coefficients (R), calculated from whole points, are low in both of the correlations.

**Supplementary Figure 5.**
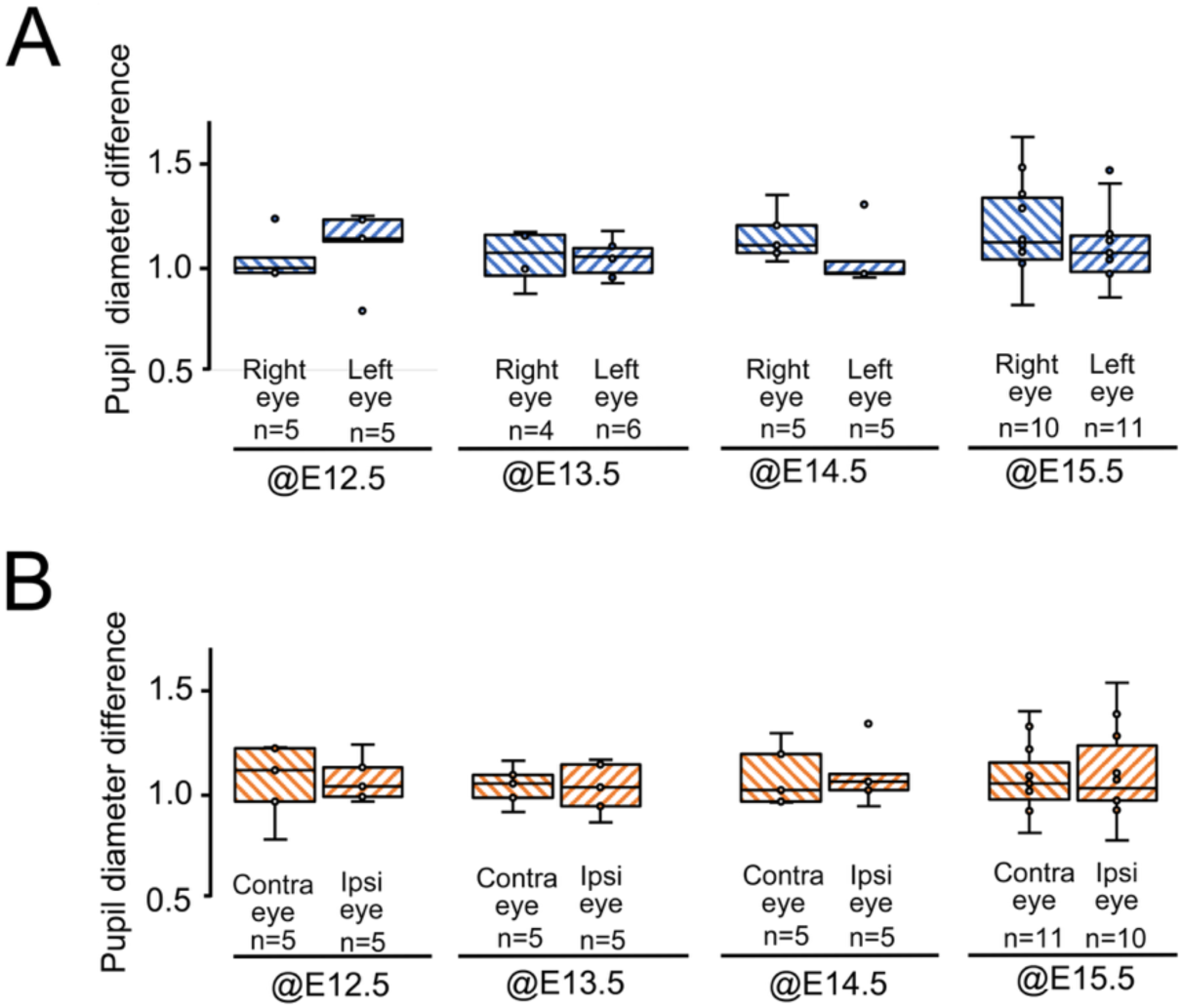
**A, B:** Boxplot showing the difference in the pupil diameter of right and left eyes (**A**) and contralateral and ipsilateral eyes to the EP side (**B**). The same mice in Fig. 5 were analyzed. Welch’s *t*-test revealed no statistical significance between the two eyes.

